# Evolutionary dynamics of pro-inflammatory caspases in primates and rodents

**DOI:** 10.1101/2024.06.19.599744

**Authors:** Mische Holland, Rachel Rutkowski, Tera C. Levin

**Author notes:** Address correspondence to Tera C. Levin. No conflicts of interest to declare.

## Abstract

Caspase-1 and related proteases are key players in inflammation and innate immunity. Here, we characterize the evolutionary history of caspase-1 and its close relatives across 19 primates and 21 rodents, focusing on differences that may cause discrepancies between humans and animal studies. While caspase-1 has been retained in all these taxa, other members of the caspase-1 subfamily (caspase-4, -5, -11, -12, and CARD16, 17, and 18) each have unique evolutionary trajectories. Caspase-4 is found across simian primates, whereas we identified multiple pseudogenization and gene loss events in caspase-5, caspase-11, and the CARDs. Because caspases-4 and -11 are both key players in the non-canonical inflammasome pathway, we expected that these proteins would be likely to evolve rapidly. Instead, we found that these two proteins are largely conserved, whereas caspase-4’s close paralog, caspase-5, showed significant indications of positive selection, as did primate caspase-1. Caspase-12 is a non-functional pseudogene in humans. We find this extends across most primates, although many rodents and some primates retain an intact, and likely functional, caspase-12. In mouse laboratory lines, we found that 50% of common strains carry non-synonymous variants that may impact the functions of caspase-11 and -12, and therefore recommend specific strains to be used (and avoided). Finally, unlike rodents, primate caspases have undergone repeated rounds of gene conversion, duplication, and loss leading to a highly dynamic pro-inflammatory caspase repertoire. Thus we uncovered many differences in the evolution of primate and rodent pro-inflammatory caspases, and discuss the potential implications of this history for caspase gene functions.

## Introduction

Pathogen-interacting genes are among the most rapidly evolving in mammalian genomes [1]. Because of the strong selective pressures involved, such genes often diversify through evolutionary arms races, which involve multiple, recurrent changes in host and pathogen molecules as each protein reciprocally adapts [2]. Common mechanisms of diversification in arms races include gene family expansion and contraction, gene conversion, and rapid amino acid turnover, particularly at sites of host-pathogen binding [2–4].

Innate immune pathways, including those involved in inflammation, are key defenses against many human pathogens and deficiencies in these pathways lead to heightened pathogen susceptibility (e.g. [5]). Over-activation of inflammation can also be dangerous for the host, both during infection [6] and in autoimmunity [7]. Therefore, we expect natural selection to act strongly on genes in inflammatory pathways, tuning these proteins to recognize an ever-changing array of microbial infections while avoiding self-harm. Indeed, some of the proteins involved in these pathways (e.g. inflammasome proteins NLRPs & CARD8) have been found to rapidly diversify, with signatures of evolutionary arms races [8–10]. We therefore sought to examine the evolutionary history and diversity of the pro-inflammatory caspase genes, which participate in these pathways.

Caspase-1 is a protease that is a hub of pro-inflammatory signaling. In the ‘canonical’ signaling pathway, multi-protein complexes called inflammasomes assemble in response to cellular or microbial signals [11]. Caspase-1 is recruited to inflammasomes via its CARD (caspase activation and recruitment domain), where its oligomerization leads to self-cleavage and activation. The active caspase-1 protease (made up of p20 and p10 subunits, see Fig. 4 for domain architectures) then targets for maturation the pro-inflammatory cytokines IL-1beta and IL-18, as well as gasdermin D, which forms membrane pores and leads to an explosive cell death known as pyroptosis (Xu et al. 2024; Bibo-Verdugo and Salvesen 2024).

In addition, there are several caspase-1 homologs (collectively known as the caspase-1 subfamily), which all reside within the same syntenic locus as the caspase-1 gene in primates and rodents [12] (Fig. 1A). One of these genes, rodent *Casp11*, has two homologs in humans, called *CASP4* and *5*, which have been implicated in ‘non-canonical’ inflammatory signaling. (Note: although rodent *Casp11* is clearly the ortholog of primate *CASP4/5* and is sometimes called *Casp4*, we refer to it here as *Casp11* as this name is more commonly used in the biomedical, mouse-centric literature.) In the non-canonical pathway, caspase-4 or -11 directly binds to cytosolic, bacterial LPS, leading to caspase oligomerization, activation, cleavage of cytokines and gasdermin D, and inflammatory cell death [13,14]. Thus, the non-canonical pathway can sense and respond to cytosolic bacterial pathogens through direct caspase activation. Some caspase-5 mutations have been associated with cancer and it is largely assumed that it participates in similar non-canonical signaling[14]. However, the molecular functions of caspase-5 have remained more enigmatic, with mainly *in vitro* evidence supporting similar biochemical activities to its caspase-4 paralog(Shi et al. 2014; Eckhart and Fischer 2024).

**Figure 1.**
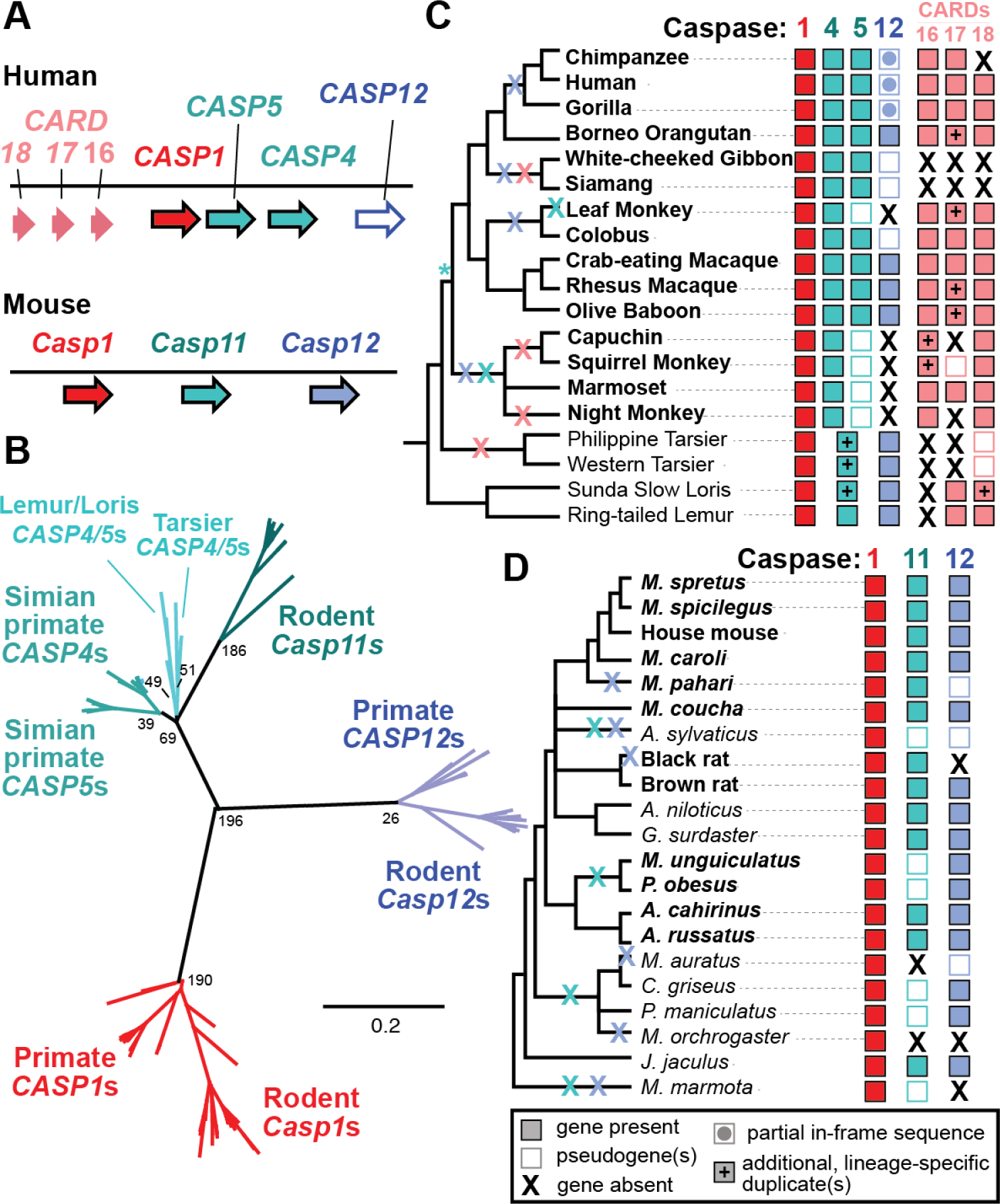
Differential patterns of gene gain and loss within the caspase-1 locus. A) Genes in the caspase-1 locus in human and mouse (not to scale). B) PhyML phylogenetic tree of intact caspase genes demonstrates that orthologs have been correctly categorized, despite gene gains and losses. Numbers at nodes are aLRT support values. C-D) Gene presence/absence across primate (C) and rodent (D) species, with colored Xs showing the history of gene losses, while * marks the *CASP4/5* duplication event. See text for definitions of pseudogene vs. partial sequence vs. gene absence. *CASP1* has been retained across all species examined, as has *CASP4* in simian primates. In contrast, there have been multiple cases of gene degradation and loss of rodent *Casp11*, primate *CASP5*, and the primate *CARD* genes. Early-branching primates encode homologs of a *CASP4/5* ancestor, with additional duplications in some lineages. *Casp12* is retained and functional across most (but not all) rodents, while it has been lost from most primates. Species in bold were used in selection analyses.

Other caspase-1 subfamily genes with less characterized roles include caspase-12, which is generally considered non-functional in humans [15–17], and three *CASP1*-like genes that contain a CARD as their only protein domain (the CARDs, aka CARD-only proteins, or COPs). The CARD-only genes are named *CARD16, 17*, and *18* (or alternatively *COP1, Inca*, and *Iceberg*, respectively) [18–20]. Overexpression studies of the CARDs suggested that they may act as caspase-1 inhibitors [18,19,21] and *in vitro* studies suggested that CARD16 and 18 promote CASP1 CARD oligomerization whereas CARD17 inhibits it [22]. However, a recent *in vivo* study of CARDs found that they each inhibit caspase-1 activation by competitively binding to the CARDs of caspase-1 and inflammasome adapters, blocking caspase-1 recruitment to inflammasomes [23]. These CARDs apparently act non-redundantly, dampening caspase-1 activation during different stages of inflammasome signaling.

In terms of their expression patterns, mouse *Casp1*, *Casp11*, and *Casp12* all have similar qualitative expression patterns to human *CASP1* and *CASP4*, i.e.) relatively high expression across most tissues and elevated in blood, spleen, and lung (Ringwald et al. 2022) (human data from https://www.gtexportal.org/ on 8/15/24). *CARD16* has a similar expression pattern, whereas *CARD17* and *18* are expressed at very low levels. The basal expression of *CASP5* is also very low except in blood, colon, and small intestine, although it has been reported that *CASP5* expression is induced by LPS and interferon-gamma (Lin et al. 2000).

Pathogens have abundant mechanisms to evade and shut down these inflammatory signaling pathways, including antagonists that act both upstream [24,25] and downstream [26] of caspase-1 activation, as well as multi-caspase inhibitors that can target caspase-1, -4, and -5 by forming covalent bonds at the caspase active site [27,28]. Specific microbial effectors that bind caspase-4 and -11 have also been identified, including the OspC3 effector from *Shigella* that inhibits caspase function through ADP-riboxination[29]. Because the members of the caspase-1 subfamily are key players in immune defenses, which can directly bind pathogen molecules and are often antagonized by pathogens, we hypothesized that they may evolve in evolutionary arms races, similar to those that have diversified inflammasome proteins[30].

Previous studies have identified some hints that the caspase-1 subfamily has been evolutionarily dynamic. For example, instances of gene duplication and loss have been observed in mammals, molluscs, and insects[12,31]. Some of the diversity of caspases has interesting and important functional implications, including the caspase-1/4 fusion proteins in carnivores[32]. Because mice are the most commonly used animal models of pro-inflammatory signaling pathways, it is particularly important to understand the similarities and differences in caspases between primates and rodents. However, prior studies of caspase evolution included at most 4 distantly related rodents and 3 primates[12], limiting our evolutionary resolution in these clades.

Here, we take an in-depth comparison of evolution of the inflammatory caspase locus across 19 primate and 21 rodent species. We find that the caspase-1 locus is much more evolutionarily dynamic in primates than in rodents, with evidence of recurrent gene duplications, pseudogenization, and gene conversion events. Contrary to our expectations, the signatures of selection were strongest and most evident in primate *CASP1* and *5* as compared to the other caspases, with interesting phenotypic implications. We also found that many commonly used mouse strains carry one or more non-synonymous variants in caspase-11 and -12, which may contribute to variation in inflammatory responses across animal studies. Combined with prior evidence of human segregating polymorphisms[33] and copy number variation[34] within the caspase-1 locus, we propose that these genes have not only a history of diversification, but also abundant, ongoing genetic innovation.

## Results

### Differential caspase gene birth and loss in primates and rodents

Caspase-1 subfamily genes all reside within a single syntenic locus (Fig. 1A), found in humans on chromosome 11q22.3. To assess the repertoires of these genes across species, we analyzed the locus across 19 primate and 21 rodent genomes. These species were selected based on: (1) sampling taxa across the phylogeny that were not too diverged from our reference human and house mouse species [i.e. *CASP1* dS<0.3], and (2) whether the full, ∼350kb locus was well-assembled on a single contig in the genome. Within the locus, we identified caspase genes by searching for exons that had 60% nucleotide identity to the annotated human or house mouse genes. In cases where diverged exons were not identified, we used synteny and alignments to the reference coding sequences to locate the missing exons.

Through this process, we classified every homolog into one of four categories. First, we called genes as “present” if they were found within their syntenic location and had a conserved, in-frame sequence that aligned across its full length to the reference. If our initial searches failed to find the homolog or identified only partial genes, we used blastn to search the Refseq Representative Genomes database for the annotated transcripts for each homolog. If this search yielded evidence of an in-frame transcript, we then identified each exon in the locus, realigned the coding sequence based on transcript evidence, and called the gene as “present”. Some homologs only had shorter in-frame sequences that did not span the full length of the reference gene. We categorized these homologs in the second category, “partial in-frame sequence”. If we identified some traces of the gene but did not find evidence of an in-frame coding sequence in either the genome or transcript database, we placed them in a third category, “pseudogenes”. Finally, if the gene was undetectable and/or if we found fewer than 70% of the exons as compared to the reference gene, we categorized the homolog as “absent”.

For all caspases that were present and intact, we confirmed their identities using a phylogenetic tree, which showed that all genes were correctly categorized into their *CASP1, 4*, *5, 11*, or *12* gene families (Fig 1B). As expected based on prior studies [12,31], this analysis also showed that *CASP4* and *5* are paralogs found only in primates and most closely related to the rodent *Casp11* gene. Yet, when we assessed which species had which intact caspase homologs to understand the history of gene gains and losses, we uncovered some surprises as well.

### Caspase-1

*CASP1* was present within the syntenic locus and we recovered full-length, well-aligned coding sequences for all 40 primate and rodent species evaluated (Fig. 1C, 1D). This pattern of widespread gene retention implies that *CASP1* plays an important, non-redundant role in organismal fitness across primates and rodents.

### Caspase-12

For *CASP12*, we found that the previously reported partial gene loss in humans [15,16] extended across most primates, with multiple lineages experiencing gene degradation over time. Importantly, we did find intact *CASP12* in the orangutan, Old World Monkeys, loris, lemur, and tarsier genomes. These are the first reported cases of intact *CASP12* in primates. Their distribution across the tree implies that *CASP12* has experienced at least four gene loss events across primate history (Fig 1C). Neither the SHS>SHG mutation found within humans that disrupts catalytic activity [15] nor the residue 125 human stop codon polymorphism[17] was shared between humans and other primates, although many primate species had numerous *CASP12* frameshifts and early stop codons. In rodents *Casp12* was present in most species.

However, we estimate there have been up to six additional instances of independent, lineage-specific *Casp12* pseudogenization and/or loss events within rodents (Fig. 1D), suggesting that *Casp12* loss is frequent across multiple mammalian clades. To test if CASP12 is slowly becoming a pseudogene across rodents as it has in primates, we made an alignment of intact rodent *Casp12* sequences and analyzed it with CODEML (model 0 vs. 0a)[35]. We detected strong evidence that rodent *Casp12* is under purifying selection (Supp Table 3), suggesting that it is likely a functional gene in most rodent species. We were not able to perform this analysis for primate *CASP12*, as we did not recover enough *CASP12* homologs at the appropriate phylogenetic distance.

### Caspase-4, -5, and 11

For *CASP4, 5*, and *11*, because these proteins are thought to share similar immune functions, we thought they might exhibit similar patterns of evolution.

Alternatively, because recently duplicated genes often lose one paralog and return to single copy, we expected *Casp11* might be conserved across rodents, whereas *CASP4* and/or *CASP5* would experience some reciprocal losses in primates. In contrast to both hypotheses, we found *CASP4* was present and shared across all simian primate genomes whereas there were multiple cases of pseudogenization (and some cases of complete gene loss) in rodent *Casp11* and simian primate *CASP5* (Fig. 1C & D). Because of these losses, four diverse rodent species, *A. sylvaticus, M. auratus, M. orchrogaster, and M. marmota,* have *CASP1* as their only caspase-1 subfamily gene.

Interestingly, in basal-branching primates (tarsier, loris, and lemur), we found that these species did not encode *CASP4* or *CASP5*, but rather a homolog of the *CASP4/CASP5* ancestor, which we here name the *CASP4/5* genes (Fig. 1 B&C). We can thus date the caspase-4/5 duplication event as occurring between 43 - 69 million years ago, in the ancestor of simian primates (Kumar et al. 2017). Notably, this conclusion differs from prior studies that have not sampled primates as thoroughly as the analysis presented here (Eckhart and Fischer 2024). In the tarsiers and Sunda Slow Loris, we discovered that the *CASP4/5* genes had undergone additional, lineage-specific duplications, independent of the duplication event that birthed *CASP4* and *CASP5*.

### *CARD* genes

Within primates, we also found at least four species that lack all three *CARDs* (Fig 1C). Based on the distribution of *CARDs* across the primate tree, we can infer that *CARD17* & *18* originated in the last common ancestor of all primates, ∼74 million years ago[36]. However, since then all three *CARDs* have experienced multiple, repeated instances of gene loss.

### Recent, repeated gene conversion and duplications in the primate *CASP1* locus

In addition to the caspase and CARD genes described above, the human locus contains multiple caspase-like pseudogenes, indicating that there have been additional historical duplication events followed by gene loss. These include two *CASP1*-like pseudogenes that lie between the *CARDs* (*Casp1P1* and *P2*) and a *CASP4*-like pseudogene (*Casp4LP*) near *CASP12* (Fig. 2). Inspired by this history of innovation, we next examined the nucleotide sequence of the locus for evidence of recent duplications or gene conversion events that may not be reflected in simple gene presence/absence data. To do so, we searched for extended blocks of similar nucleotide sequence, defining blocks of self-similarity as regions at least 1kb long that contained at least 50% nucleotide identity to another region within the 350kb locus.

**Figure 2.**
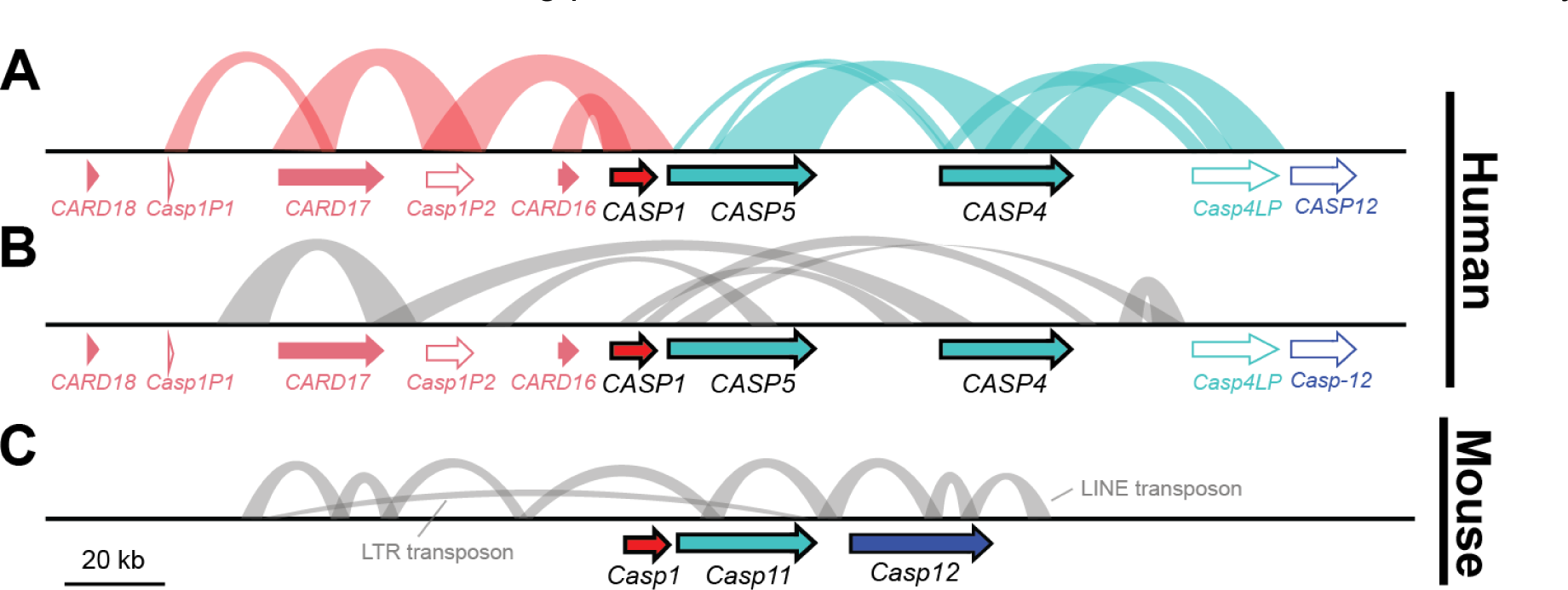
The caspase-1 locus has undergone recent gene conversions and tandem duplications in primates, but not rodents. A-C) Annotated genes (filled colored arrows) and pseudogenes (white arrows with colored outlines) in the human (A-B) and mouse (C) loci. Colored arcs (A) connect regions of extended nucleotide identity between caspase genes or pseudogenes, indicative of recent tandem gene duplications or gene conversions. Gray arcs (B and C) show regions of sequence identity within intronic or intergenic regions.

In the human locus, we detected many regions of self-similarity, including those that duplicated both coding and non-coding sequences (Fig. 2A & B). For example, a ∼6kb region of the *CASP4* locus (including both exons and introns) was 60% identical to the *CASP4LP* pseudogene and 54% identical to the *CASP5* locus. We found *CASP4LP* loci only within great apes (and it was a pseudogene in each case), suggesting that this locus likely arose ∼13Mya, whereas CASP5 dates back to the last common ancestor of simian primates ∼50 Mya[36] (Fig 1). These findings (and those of Fig 3 & 4 below) suggest that *CASP4* and *CASP5* have experienced recent gene conversion events that overwrote the sequence of one locus with the other at one or more times during primate evolution. We found similar blocks of self-similarity in the human locus between *CASP1* and *CARD16, CARD17*, and nearby pseudogenes (Fig 2A), as well as blocks that included intronic, intergenic, or promoter regions (Fig 2B). Although *CARD18* arose via *CASP1* duplication at approximately the same time as *CARD16* & *17* (Fig. 1), it appears that *CARD18* has not undergone recent gene conversion, as *CARD18*’s sequence is now too diverged to be identified in this analysis. Overall, recurrent segmental duplications and gene conversion events within the primate CASP1 locus appear to be common.

**Figure 3.**
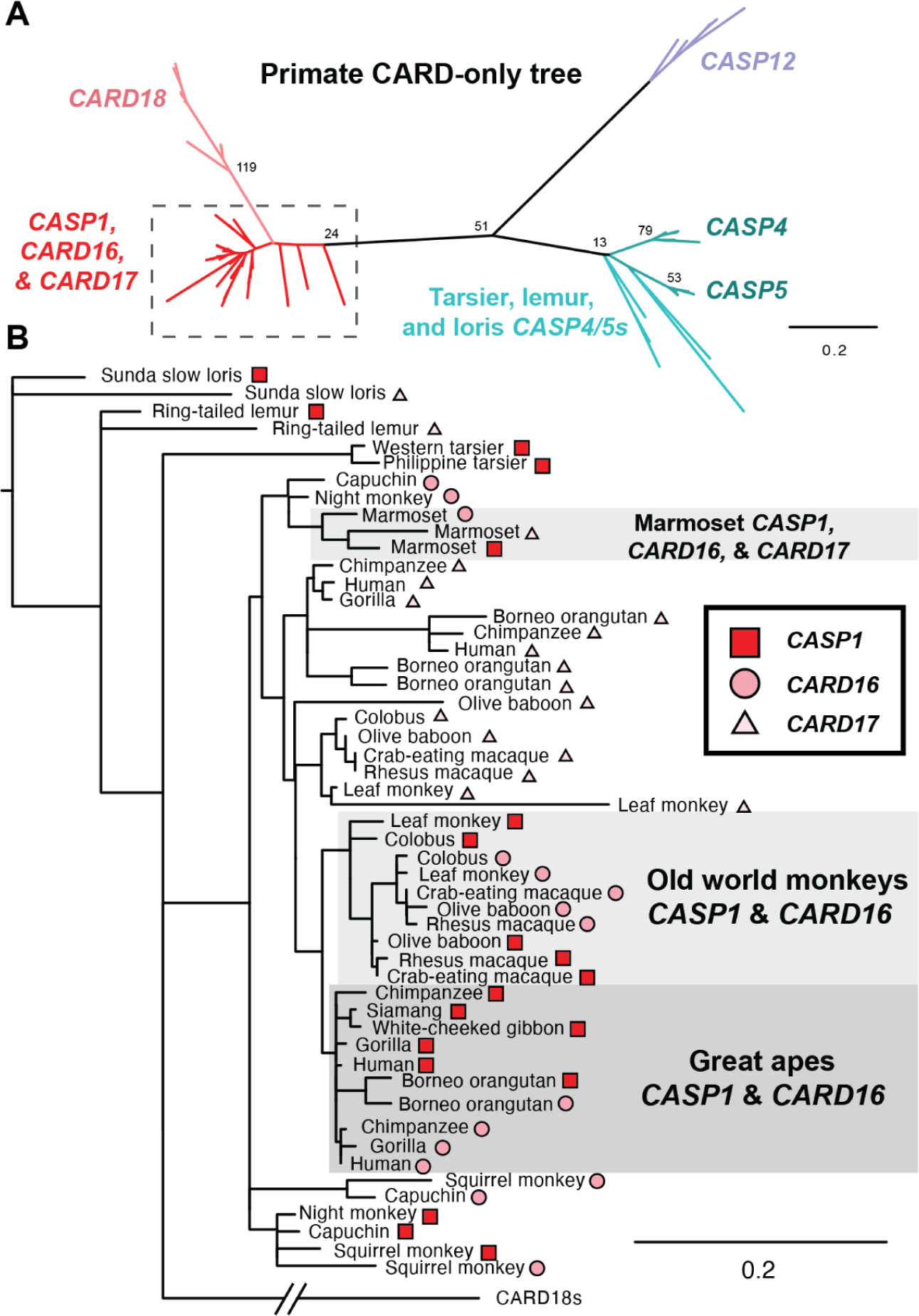
PhyML phylogenetic tree of the CARDs of each primate caspase-1 subfamily gene. Region within the dashed box in A) is shown in detail in panel B), with gene identity indicated by symbols. Although sequences for *CASP4/5s*, *CASP12,* and *CARD18* have distinct branches, in several species the *CASP1*, *CARD16*, and/or *CARD17* sequences are highly similar and intermixed on the tree, indicating recent gene conversion. All illustrated nodes on the PhyML trees had aLRT support statistics >5. Other nodes were collapsed.

These events have significantly impacted the sequences, and potentially functions, of genes in the primate locus, as illustrated by a phylogenetic tree of primate CARD sequences (Fig 3A). In this tree, we observed that the *CASP4/5, CASP12,* and *CARD18* branches were clearly differentiated from each other, whereas there was an intermingling of the *CASP1*, *CARD16*, and *CARD17* sequences (Fig 3B). Specifically, we saw that the sequences of *CASP1* and *CARD16* were intermixed together in Old world monkeys, independently intermixed within Great apes, and that the sequence of marmoset *CASP1* was most similar to marmoset *CARD16* and *17*. Based on gene synteny and the presence of *CARD16* and *17* across primates, we therefore infer that these genes have undergone recent gene conversion with *CASP1* in at least three instances. Because these events overwrite the *CARD* sequence with that of *CASP1*, we relied on synteny to define the presence and absence of *CARD16* and *17* in Fig 1. We predict that such gene conversion events would enable CARD16 and CARD17 to retain strong, homotypic binding to the CASP1 CARD, despite divergence in the CASP1 CARD sequence over time.

In contrast to the highly dynamic human locus, few blocks of self-similarity exist within the house mouse locus (Fig. 2C). Indeed, we did not detect any instances of extended nucleotide identity that included coding regions of any caspase. We did detect multiple transposon insertions within intergenic and intronic regions (specifically the Long interspersed nuclear element L1MdMus and ERVL-MaLR family of LTR transposon [37]), demonstrating that we had the power to detect such duplication events. We also performed similar phylogenetic analyses in rodent genomes as in Fig 3, but detected no evidence of gene conversion between rodent *Casp1, Casp11*, or *Casp12.* Thus, unlike in the human locus, we did not find any evidence in the mouse genome of ongoing caspase-1 subfamily duplication or gene conversion events.

Finally, within the human population it appears that duplications within the *CASP1* locus are ongoing. Copy number variation among humans is common, including between the GRCh38 genome assembly, which we focused on here, and the recent telomere-to-telomere genome sequence of the CHM13 cell line, which derives from a single human haplotype (T2T-CHM13) [34].

While segmental duplications tend to get collapsed in most genome assemblies, the T2T-CHM13 genome is currently unique in its completeness and ability to capture structural variation. In this haploid genome, the *CASP1* locus was identified as one of the regions with highly variable gene dosage, with T2T-CHM13 containing an estimated one copy of *CARD16*, two copies of *CASP12, CARD 17*, and *CARD18*, three copies of *CASP1*, five copies of *CASP5*, and fourteen copies of *CASP4*. Therefore, throughout primate history and within humans, the CASP1 locus appears to be a frequent site of gene duplication and innovation.

### Unexpected patterns of positive selection in the caspase-1 subfamily genes

Canonical pro-inflammatory pathways activate caspase-1 downstream of pathogen recognition. In contrast, caspase-4 and -11 have been found to directly bind multiple types of pathogen molecules [13,29,38], whereas no pathogen antagonists have yet been identified that specifically target caspase-1. Because both LPS and microbial effectors tend to be highly variable and because we expect to see rapid evolution at host-pathogen binding interfaces, we hypothesized that we would see strong signatures of positive selection within *CASP4* and *Casp11*, potentially in the previously-defined LPS-binding residues of the CARD region[13], whereas these signatures might be weaker or absent in *CASP1* and *CASP5*.

To test this hypothesis, we analyzed evolutionary genomic signatures in alignments of primate *CASP1, CASP4*, and *CASP5* as well as rodent *Casp1* and *Casp11*. Recurrent gene conversion events (as discussed above) can cause different parts of a gene to have different evolutionary histories. To detect if this was the case and identify sites of recombination breakpoints, we used the program GARD [39]. Consistent with our earlier results, GARD identified multiple recombination breakpoints across the caspases, with more sites identified in the primate than in the rodent genes: two recombination breakpoints in primate *CASP1*, one breakpoint in *CASP4*, three breakpoints in *CASP5*, one breakpoint in rodent *Casp1*, and none in rodent *Casp11*. Thus, many extant caspase genes are evolutionary patchworks, reflecting multiple gene conversion events.

This sort of recombination history can interfere with tests for positive selection. Therefore, we divided each alignment into different recombination segments and analyzed each segment independently. For each, we used multiple algorithms to estimate the dN/dS ratio, i.e. the rate of non-synonymous to synonymous substitutions. If the dN/dS ratio is significantly higher than 1, that would indicate that the protein regions and amino acid sites have evolved under positive selection, with more rapid amino acid changes than expected by neutral mutation. For these statistical tests of positive selection, we used CODEML and FUBAR [35,40]. Both algorithms are designed to detect positive selection that is pervasive across the phylogenetic tree and they use different statistical models of molecular evolution. Therefore, these analyses detect related, but complementary, signatures of positive selection.

Contrary to our initial hypothesis, CODEML did not detect significant evidence of positive selection in primate *CASP4* or rodent *Casp11* (Table 1, Fig 4A & B). Similarly, for these genes only a few sites were called by FUBAR: two sites in the CARD of *CASP4* and two in the p20 subunit of *Casp11*. None of these sites overlapped with the LPS-binding residues proposed by Shi et al., 2014[13]. Of the FUBAR-identified sites, the most convincing was I156 in the p20 domain of mouse *Casp11*, which toggled between I, S, T, and N codons across the 10 rodents analyzed. Interestingly, I156 was also the only residue where homologous sites in primate *CASP1* and *CASP5* were also identified as positively selected (see below). In addition, this site is in proximity to the binding interface between caspase-4 and -11 to a bacterial effector from *Shigella*[29], suggesting that this site may evolve rapidly to evade effector binding. However, with the possible exception of the I156 residue, we found little to no evidence that *CASP4* or *11* have evolved under positive selection, with no enrichment at putative LPS binding sites.

**Figure 4.**
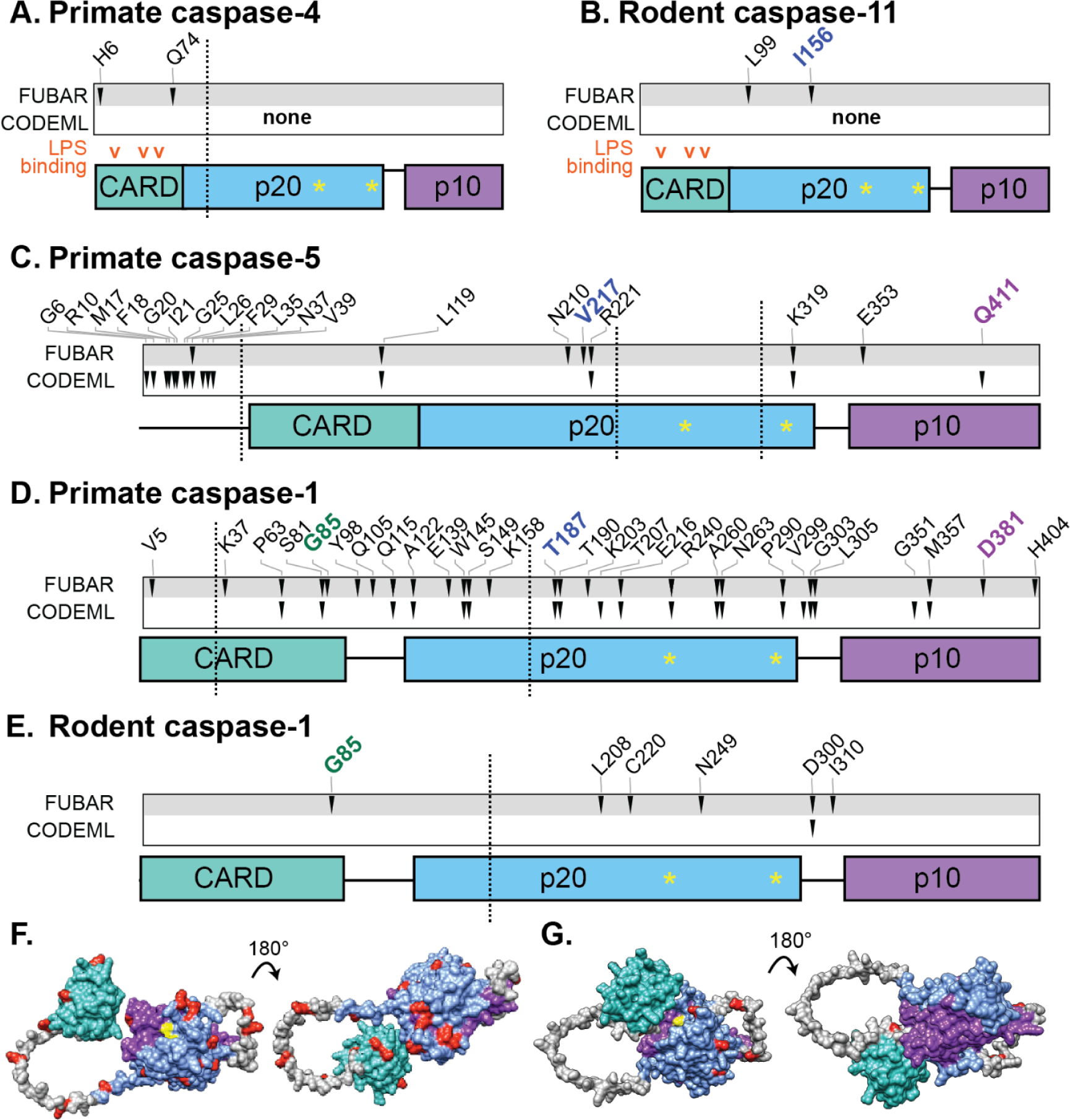
Positively selected residues in pro-inflammatory caspases in primates and rodents. A-E) Sites of positive selection, as determined by the FUBAR and CODEML algorithms. Arrowheads mark sites passing the statistical threshold for positive selection. Vertical dotted lines separate recombination segments identified by GARD, which were each analyzed independently. The domain architecture of each caspase is shown as colored boxes, with active sites illustrated (yellow *) along with the proposed LPS-binding residues of CASP4 and 11 according to Shi et. al 2014 (orange carats). In three cases, sites of positive selection in primate CASP1 were homologous to sites identified in one or more other caspase alignment (green, blue, or purple residue names). All coordinates are named according to the residues in the human or mouse sequences for primate or rodent analyses, respectively. F-G) Each identified site of CASP1 positive selection (red) is illustrated on the predicted AlphaFold structure of human (F) or mouse (G) caspase-1. The CARD, p20, and p10 domains are colored as in (A-E), with linker regions in gray, and active sites in yellow. Many more sites of positive selection were identified in primates than in rodents, but in neither clade do the sites show much co-localization in 3D space.

**Table 1.** Signatures of selection in caspase genes.

| Gene | Clade | Recomb. segment | dN/dS | M7 v. M8 p-value | M8 v. M8a p-value | Highest omega in M8 | # of positively selected sites in CODEML | # of positively selected sites in FUBAR |
| --- | --- | --- | --- | --- | --- | --- | --- | --- |
| *CASP1* | Primate | 1 | 0.41516 | 9.31E-01 | 9.44E-01 | 3.60 | 0 | 1 |
|  |  | **2** | **1.36624** | **8.93E-09** | **1.00E-08** | **9.56** | **6** | **12** |
|  |  | **3** | **0.68748** | **6.51E-07** | **6.54E-07** | **3.72** | **13** | **14** |
| *CASP4* | Primate | 1 | 0.35825 | 5.01E-01 | 6.04E-01 | 2.43 | 0 | 2 |
|  |  | 2 | 0.23668 | 1.40E-01 | 1.75E-01 | 6.96 | 0 | 0 |
| *CASP5* | Primate | **1** | **3.15359** | **2.90E-03** | **2.91E-03** | **5.01** | **12** | **1** |
|  |  | 2 | 0.99276 | 1.80E-02 | 1.85E-02 | 2.33 | 2 | 4 |
|  |  | 3 | 0.3462 | 9.96E-01 | 9.73E-01 | 1.00 | 0 | 0 |
|  |  | 4 | 1.09378 | 1.60E-02 | 1.61E-02 | 5.27 | 2 | 2 |
| *Casp1* | Rodent | 1 | 0.58534 | 3.60E-01 | 6.18E-01 | 1.29 | 0 | 1 |
|  |  | 2 | 0.33065 | 8.47E-02 | 1.61E-01 | 1.83 | 1 | 6 |
| *Casp11* | Rodent | 1 | 0.41356 | 3.65E-01 | 7.32E-01 | 1.37 | 0 | 2 |
| *Casp12* | Rodent | 1 | 0.19438 | 1.00E+00 | 1.00E+00 | 24.07 | 0 | 0 |
|  |  | 2 | 0.51008 | 1.00E+00 | 1.00E+00 | 1.00 | 0 | 0 |
|  |  | 3 | 0.17643 | 1.00E+00 | 1.00E+00 | 1.00 | 0 | 2 |

In contrast, we identified significant signatures of positive selection in both primate *CASP1* and *CASP5.* For *CASP5*, we analyzed sequences only in apes and Old world monkeys, because *CASP5* in New world monkeys appears to be a pseudogene (Fig 4). Nevertheless, despite the reduced statistical power from fewer sequences, we identified 19 sites under positive selection in *CASP5*, 4 of which were called by both programs (Fig 4C). Unlike *CASP4*, *CASP5* has an extended N-terminus. This N-terminal region was enriched for positive selection hits (56% of total sites identified by both algorithms were located within the first 12% of the coding sequence length), with CODEML calling an especially large number of sites. In CODEML, positive selection testing can be less powerful in short alignments. Therefore, it is possible that we have missed some of the positively selected sites on this short, N-terminal segment. The *CASP5* N-terminal tail also included an insertion within Old world monkeys, further diversifying this region. When we analyzed the N-terminal region both before and after trimming out the Old world monkey insertion, CODEML and FUBAR identified many positively selected sites in both cases, although the identities of the sites shifted. Therefore, while we are confident that the N-terminal region of *CASP5* is rapidly evolving, the specific sites identified should be interpreted with caution. While the N-terminus of *CASP5* has not previously been associated with a known function, the diversification observed here could indicate that this part of the protein is important for organismal fitness. We also note that *CASP5* has multiple annotated splice isoforms, some of which exclude exon 2 which encodes most of the N-terminus. Therefore, although these residues passed our statistical thresholds in one or more algorithms, it’s possible that the elevated rate of amino acid turnover in the N-terminus is due in part to relaxed selection, as this region is less frequently part of the caspase-5 protein.

Even ignoring the *CASP5* N-terminus, we still detected more residues under positive selection and with stronger support in *CASP5* than in its close paralog *CASP4* or rodent *Casp11* (Fig 4; Table 1). Of particular note, three positively selected sites were homologous between primate CASP5 and CASP1, including V217, homolog of I156 in caspase-11. Within humans, the V217 residue of CASP5 is polymorphic at high frequencies (11% of individuals in Africa and 4% worldwide have L or M at this site) [33], demonstrating that variation at this site still segregates within human populations, not only across different primate species.

Finally, in *CASP1*, we identified 29 sites of positive selection primates (16 called by both CODEML and FUBAR) and 6 in rodents (1 called by both algorithms (Fig. 4D & E)). These sites were distributed across the length of the protein, including sites within the CARD, p20, p10 domains, and the flexible linkers connecting these domains. Positively selected residues that are distant in the primary sequence of a protein will sometimes co-localize in the folded protein, often overlapping with host-pathogen binding interfaces. To see if the *CASP1* selected residues potentially formed this kind of interface, we examined them on the AlphaFold predicted structures of primate and rodent caspase-1, but did not observe a large concentration of the residues in 3D space. The localization was similar when we used experimental caspase-1 crystal structures (not shown).

In summary, we detected little to no evidence of positive selection in *CASP4* and *Casp11*, whereas there were strong signatures of selection in primate *CASP5* and *CASP1*. In three cases, sites we identified in primate *CASP1* were homologous to positively selected sites in other caspases. However, with the exception of the N-terminal tail of *CASP5*, the positively selected sites were not concentrated or co-localized in a particular region of the protein.

### Variation in caspase-1 aspartate self-cleavage sites

During caspase protease activation, protein oligomerization induces self-cleavage at aspartate sites to generate the active enzyme [41]. The removal of the flexible linker between the p20 and p10 subunits is essential for caspase-1 activation. The aspartate cleavage sites flanking this linker are strictly conserved across caspase-1, 4, 5, and 11 (Fig 5). However, it has been open to debate whether the CARD-p20 cleavage is essential for pro-inflammatory caspase function. When we examined the aspartate site at the N-terminal end of p20, we found it was conserved across caspase-4, 5, and 11, with the exception of one rodent species, *M. pahari*, which had an asparagine at the caspase-11 site instead. Across caspase-1 homologs, we found many more species that had variation in the CARD-p20 cleavage site (Fig 5), often with no aspartate residues nearby that could serve as alternate cleavage sites. These species included all of the New world monkeys and four species of rodents, including rats. We suggest that functional studies of the caspase-1 homologs from these lineages may provide important information about the mechanisms and activation of pro-inflammatory caspases, particularly the role of CARD-p20 cleavage.

**Figure 5.**
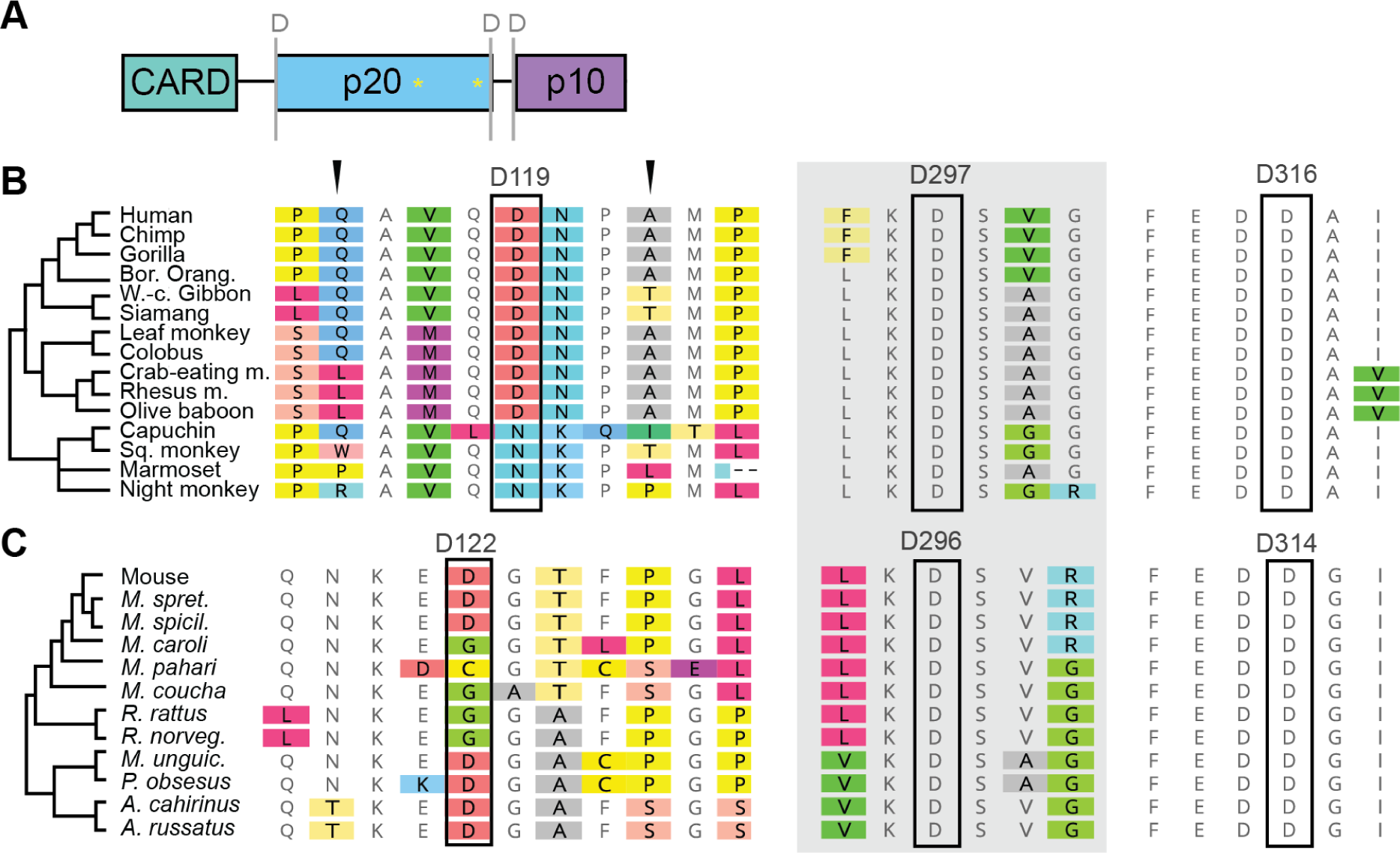
Caspase-1 aspartate cleavage sites between CARD and p20 domains are not strictly conserved. A) domain architecture of caspase-1, showing locations of aspartate cleavage sites (’D’). B-C) Alignments of the caspase-1 aspartate cleavage sites, with residues that do not match consensus highlighted with colored boxes. While the two C-terminal cleavage sites are highly conserved within primates (B) and rodents (C), the N-terminal site is more highly variable and some species lack the aspartate residue entirely. Arrowheads mark nearby positively selected sites.

### Caspase variants within inbred mouse and rat lines

While inbred mouse and rat lines are widely used for studies of inflammation and immunity, these strains can harbor polymorphisms that impact caspase activity. Indeed, the non-canonical inflammasome signaling of caspase-4/11 was only discovered in 2011 after scientists realized that the widely-used 129S1 mouse strain background carried an inactivating mutation in caspase-11 (Kayagaki et al. 2011). This mutation was a 5 bp deletion that eliminated the splice acceptor on exon 7 of caspase-11, resulting in an out-of-frame splice isoform that behaved as a caspase-11 null. Inspired by this history, we looked for non-synonymous caspase-1-family mutations in the genomes of 16 mouse lines and 8 rat lines that are commonly used for laboratory study. In addition to the 129S1 deletion described above, we identified a wide variety of missense variants. By comparing these variants to our alignments, we categorized each variant as: (1) having a potential to impact caspase function (if the polymorphism altered a site highly conserved across 10 closely related rodents), (2) unlikely to impact function (if the polymorphism introduced a residue that was shared with another rodent species), or (3) unknown. We identified only a few variants across the rat strains, most of which we predicted were unlikely to affect caspase function (Table 2). One notable feature was a 1bp “insertion” in exon 7 of caspase-12 that was present in all 8 rat genomes. Whereas the reference rat caspase-12 gene has a frameshift relative to the homologs, this 1bp “insertion” resulted in a full-length, in-frame caspase-12. Because this “variant” was in every strain and because the in-frame transcript is well-represented in RNA-seq datasets (NCBI Reference Sequence: NM_130422.2), we believe this position reflects an error in the reference rat genome assembly. Otherwise, we found very few non-synonymous variants within the rat caspase-1, -11, and 12 genes.

**Table 2.** Non-synonymous variants in rat inbred lines.

|  | *Casp1* | *Casp11* | *Casp12* |  |
| --- | --- | --- | --- | --- |
| **Strain** | **Q343R** | **V60A** | **F115L** | **1bp ins** |
| ACI_N | x | x |  | x |
| BN_SsN |  |  |  | x |
| BUF_N | x | x |  | x |
| F344_N | x | x |  | x |
| M520_N | x | x |  | x |
| MR_N | x |  | x | x |
| WKY_N | x |  | x | x |
| WN_N | x |  |  | x |

Italics and light shading indicate sites that are unlikely to disrupt function, i.e. the same residue is present at that site in other rodent homologs. Dark gray boxes indicate sites with the potential to disrupt function, as the variant alters a highly conserved site. White boxes indicate the presence of the reference allele.

In contrast, we identified many caspase-11 and -12 variants in the inbred mouse lines, such that 50% of the mouse strains we examined carried missense variants in one or both genes (shaded strains, Table 3). Only the 129S1 strain carried the previously-characterized 5bp deleterious deletion in caspase-11. While the remaining missense variants will need to be tested experimentally, we predict that several are likely to impact caspase function. We therefore recommend that researchers studying caspase-11 or -12 may want to avoid using the following mouse lines: 129S1/SvImJ, cAST/EiJ, pWK/PhJ, wSB/EiJ, aKR/J, cBA/J, nOD/ShiLtJ, and IP/J. The sPRET strain contained multiple non-synonymous mutations, although we predict these are likely to have little impact. There were also 7 mouse strains that did not carry any missense variants in caspase-1, -11, or -12, which would be good models to use for caspase-11 and -12 studies: bALB/cJ, c57BL/6NJ, a/J, nZO/HILtJ, fVB/NJ, dBA/2B, and c3H/HeJ.

**Table 3.**
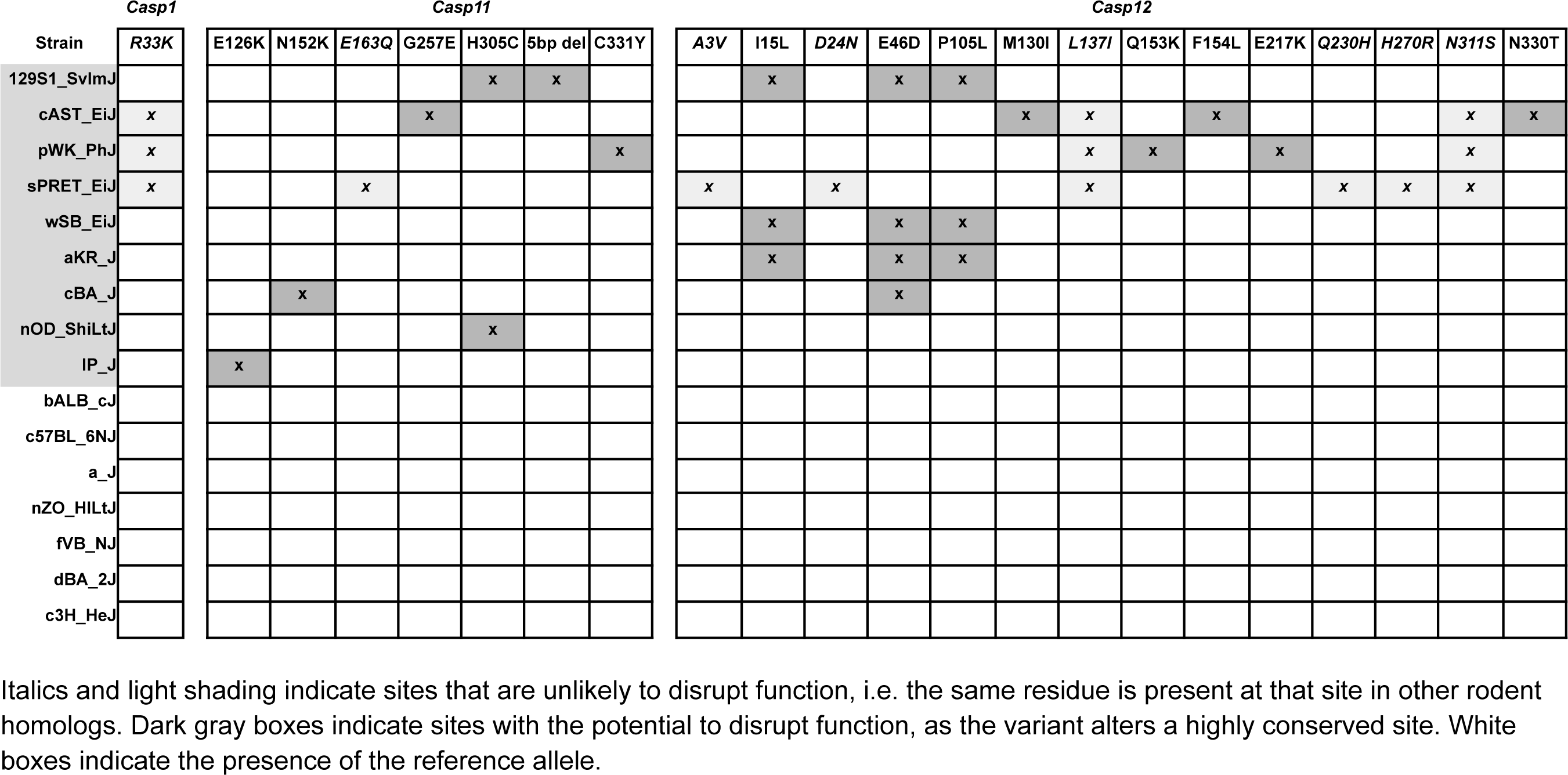
Non-synonymous variants in mouse inbred lines.

## Discussion

While researchers commonly use mice as model systems for studying human pro-inflammatory caspases, we find a number of differences in the genes and evolutionary dynamics between primates and rodents, with potential implications for caspase functions.

First, the caspase-1 locus is more evolutionarily dynamic in primates, with evidence of recurrent gene duplications, pseudogenization of new genes, and gene conversion events (Fig. 1-3). Such events can create blocks of nucleotide identity within the locus that can facilitate further rearrangements or recombination, thus perpetuating complex genetic dynamics. The rapid amino acid turnover, particularly in primate *CASP5* and *CASP1* (Fig. 4), further accelerates the evolution of these genes. Combined with prior evidence of human segregating polymorphisms and copy number variation within the caspase-1 locus, we propose that these genes have not only a history of diversification, but also abundant, ongoing innovation in primates. The genetic variation in the caspases is likely to cause differences in pro-inflammatory immune responses across individual humans (and cell lines) as well as non-human primates, which won’t be reflected in mouse models. Inversely, many rodent laboratory models harbor caspase segregating variants that are not reflective of humans (Tables 2-3). Therefore, we urge researchers to carefully assess and report the caspase genotypes and gene copy number for the organisms and cell lines used in their experimental studies.

Second, our work has uncovered new information about variation in caspase gene content among primates and rodents. The *CASP4/5* duplication event occurred in the ancestor of simian primates, around 50 million years ago[36], while the tarsier and loris lineages had additional, independent duplications of this locus (Fig 1). Because duplication events often lead to subfunctionalization or neofunctionalization if the duplicates are retained for long periods of time, it is likely that there are significant differences in the functions of *CASP4/5* genes and their single copy *Casp11* rodent homolog. Similarly, the *CARD* genes arose early near the base of the primate tree, around 74 million years ago. Because each of these genes lacks 1-to-1 orthologs in rodents and primates, their human functions may not be reflected in mouse models. We also discovered that both primates and rodents encode intact, likely functional copies of *CASP12*, although this gene has been recurrently lost in both lineages. More surprisingly, we discovered multiple pseudogenization and gene loss events of both primate *CASP5* and rodent *Casp11*, including four rodent species that apparently have *Casp1* as their only caspase-1 subfamily gene. Given the proposed importance of *Casp11* for pathogen defense, we speculate that these four species may have acquired new genes or alleles that can detect cytoplasmic LPS, thus compensating for *Casp11* loss.

The evolutionary signatures also have interesting implications for the molecular functions of the caspase-1 subfamily. For example, we were surprised to see such strong selective signatures in *CASP5* as compared to its close homologs *CASP4* and *Casp11*, because current studies have yet to identify *in vivo* phenotypes or molecular functions associated with caspase-5 (nor its unique N-terminal tail), whereas both caspase-4 and -11 are known to mediate important host-microbe molecular interactions. These evolutionary discrepancies suggest that caspase-4 and -5 may have significant differences in their *in vivo* functions and that caspase-5 plays an important, undefined, role supporting organismal fitness. For example, caspase-5 may substitute for caspase-4 in certain contexts, or it may act as a molecular decoy for caspase-4 inhibitors. Although these scenarios will remain speculative until future experimental advances in the role of caspase-5, the positively-selected residues we identified in Fig. 4C are likely to be sites that mediate caspase-5-specific activities.

We were also surprised to see the widespread signatures of positive selection in primate *CASP1*, as compared to the other caspase genes. This pattern suggests that caspase-1 is under substantial selective pressure at multiple molecular interfaces, and that these pressures are largely not shared with caspase-4 or -11. (The only exception being at I156 of *Casp11*/ V217 of *CASP5*/ T187 of *CASP1*, which could interact with similar pathogen antagonists). Analogous patterns of protein evolution (i.e. many positively selected sites distributed throughout the protein) have been found in examples such as PKR and MxB, which are thought to be on the ‘defense’ of evolutionary arms races[2]. In these scenarios, proteins are evolving to evade pathogen antagonists at multiple sites throughout the protein. Although pathogens are known to antagonize the caspase-1 signaling pathway both upstream[24,25] and downstream[26] of caspase-1, we would not expect these known, indirect antagonists to drive adaptation in caspase-1 itself. Several viruses encode multi-caspase inhibitors that can target caspase-1, -4, and -5 by forming covalent bonds at the caspase active site [27,28]. However, as we did not observe an enrichment of positive selection around the caspase-1 active site (Fig. 4D-G), and as the inhibitors are not caspase-1-specific, we also do not believe that these inhibitors fully explain the patterns of caspase-1 evolution. Instead, we postulate that there are multiple, unknown pathogen proteins that specifically bind and inhibit caspase-1 at different molecular interfaces. We further predict that the residues we highlighted here are likely to mediate variation in binding specificity between caspase-1 and various antagonists as well as between a given antagonist and different members of the caspase-1 family.

## Materials and Methods

### Selection of species and genomes to analyze

To compare the evolutionary histories of primate and rodent caspases, our analyses required at least 10-15 caspase genomic loci from both the rodent and primate clades[42]. We selected genomes to use in 2022 and therefore included the latest assembly version available at the time (Supp Table 1). High quality genome assemblies were included based on the following quality metrics: low assembly fragmentation (N50>300kbp and L50<2000), and high read coverage (>40X). Rodent and primate species trees were based on previously published work (Menezes et al. 2010; Jameson et al. 2011; McBee et al. 2015; Molaro et al. 2020; Swanson et al. 2019; Côrte-Real et al. 2022) and the Zoonomia database. To approximate divergence of the species, we used CODEML model 0 [35] to calculate the pairwise dS between the *CASP1* homologs in the species of interest and either humans (for primates) or house mouse (for rodents). Species that had a pairwise dS >0.3 from the reference were considered to be outgroups.

### Extracting the caspase-1 locus from the assemblies

To identify the *CASP1* locus and *CASP1*-related homologs in each of our genome assemblies, we first created local blast databases of the collected, high-quality genomes for rodents and primates using BLAST+ v2.13.0 [47] makeblastbd. We then used the Refseq representative transcript of house mouse or human *CASP1* as a query in a blastn search on our local databases with default settings. From the output of possible *CASP1* hits in each genome, we filtered for strong exon hits using a bitscore>100. The coordinates of the hits also revealed that each *CASP1*-like gene was contained within a single genomic locus in each species, and none were located on different chromosomes. We then identified start and end coordinates of each candidate *CASP1* homolog in each assembly, based on homology to the beginning and end of the reference query. Using the coordinates of the strongest *CASP1* hit, we then extracted the *CASP1* genomic locus with excess flanking two Mbp for each assembly.

### Exon annotations, alignments, phylogenetic trees, and caspase paralog identification

To retrieve accurate *CASP* CDS sequences from the *CASP1* locus in each species, we first annotated each exon. For each of the *CASP1* homologs and *CARD*-only proteins, we chose the Refseq representative transcript from human or house mouse as our reference CDS for exon annotation. In humans *CASP12* is considered catalytically inactive and likely a pseudogene[12,15]. Because pseudogenes are undergoing degeneration of their coding sequence, annotation of conserved exons from a pseudogenic reference is challenging.

Therefore, we performed exon annotation of primate *CASP12* loci using both the Sunda slow loris and house mouse *CASP12* CDS sequences.

We searched for conserved exons of each homolog using both nucleotide similarity and synteny. To identify exons with high nucleotide identity to the reference (>60% identity to humans in primates, or to house mouse in rodents), we used the Geneious Prime v2024.0.3 “Transfer Annotations” tool to transfer exon annotations from the reference sequence to the locus of interest. We empirically found that lowering the similarity below 60% resulted in spurious exon hits that did not align with homologs or contribute to full coding sequences after alignment. While this approach identified most exons of interest, some diverged exons were missed, whereas exons similar to multiple caspase paralogs matched to multiple regions within the same locus. We therefore relied on gene synteny to assign exons to the correct *CASP* gene and identify paralogs, even if only part of the gene was present.

To identify missing exons and obtain full CDS sequences for each gene from each species, we aligned the coding sequences of each gene with the annotated genomic DNA using MAFFT[48] with automatic direction detection. We then used the exon annotations to guide the manual trimming of introns across the alignment. We then realigned the trimmed sequences by translation using Geneious Prime translation alignment tool with the MUSCLEv5[49] plugin. We confirmed the quality of our manual trimming by comparing our trimmed reference species CDSs to the publicly available reference CDSs and protein sequences. In some cases, like the Jerboa *Casp11* and *Casp12*, our manual trimming method initially failed because the intronic regions were very different sizes compared to the rest of rodent homologs. However, using predicted CDS of the Jerboa caspases and our exon annotation, we were able to recover full-length inframe CDS of these genes through manually trimming, independent of the gDNA alignment. For primate *CASP5*, Old world monkeys had an insertion within the N-terminal tail of the protein, which we manually trimmed out before further analyses. We generated all phylogenetic trees using the Geneious Prime tree builder PhyML[50] plugin with GTR model and aLRT statistics. We constructed the trees in Fig. 1 and Fig 3 with multiple different substitution models, including GTR, HKY85, and/or Blosum62. In all cases, the trees were very similar, with only minor variation in support values and branch lengths.

### Pseudogene and gene loss calls

While *CASP1* was well-conserved in all lineages, other genes in the locus were not. To confirm the accuracy of our homolog genomic extractions, especially in cases of pseudogenes and gene absence, we searched for annotated transcript evidence of each caspase homolog. We performed an NCBI blast search in NCBI Transcript Reference Sequences database (reseq_rna) with each caspase homolog CDS reference. The presence of a full-length in-frame transcript from blast results, identical in sequence to our extracted CDS, was additional evidence of a functional full-length *CASP* homolog. If there was not a full-length in-frame transcript but an alternative splice isoform from blast results was still in-frame, we called these homologs as “partial in-frame transcripts”. However, this CDS search strategy was inconclusive if *CASP* homologs did not have any transcript evidence at all, leaving open the possibilities of gene loss, pseudogenization, or lack of genome annotation.

We called events as “gene losses” if fewer than 70% of the exons were identified as described above. These partial loci were not included in subsequent gene alignments or trees. To define pseudogenes, we examined the sequences for frameshift indel mutations, exon truncations, or exon loss. Frameshift indel mutations were differentiated from genome assembly errors by examining closely related species to see if the same defect was present in multiple, independent studies/assemblies. See Supp Table 2 for a full description of the presence, absence, and pseudogene data for each gene in each species.

### Identification of regions of nucleotide identity within the caspase locus

To search for evidence of recent gene conversion or duplication, we identified regions of extended nucleotide identity consistent with recent gene conversion in the caspase locus of humans (GCF_000001405.40) and *M. musculus* (GCA_000001635.9) (Fig 2). To do so, we extracted a ∼350kb region from each genome, centered on the *CASP1* gene. We then used the Dotplot tool of Geneious Prime to detect any regions of self-similarity, which appeared as off-diagonal lines in the dotplot. We annotated any DNA regions that were at least 1kb long that contained at least 50% nucleotide identity to another region within the 350kb locus.

### Measurements of selection and recombination

To perform selection analyses, we selected *CASP* sequences that fit within at an appropriate level of divergence. 15 primate assemblies were selected for selection analyses with the most diverged species from human being marmoset at a dS of 0.13. The rodent Muridaea clade of 12 species was selected for selection analyses with the most diverged clade from house mouse being spiny mouse at a dS of 0.29. To detect recombination breakpoints, we ran GARD[39] on each homolog nucleotide alignment with default DataMonkey settings: faster run mode, universal genetic code, no site-to-site rate variation, two rate classes. Because recombination can cause false positives in selection analyses, we then split the alignments at recombination breakpoints and analyzed each recombination segment independently. To detect residues of positive selection, we ran two softwares: PAML CODEML[35] and FUBAR[40]. We performed PAML CODEML analysis with models 0, 0a, 1, 2, 7, 8, and 8a. All PAML analyses with estimated omega were initially run with a starting omega of 0.4 and the codon frequency model F3X4. We additionally re-ran all PAML analyses with a species and segment alignment tree.

Model 0 estimates a single dN/dS value across the entire gene. Model 0a fixes the starting omega to 1 to simulate a gene evolving neutrally. Model 1 partitions sites into two dN/dS estimates, dN/dS =1 or dN/dS<1, whereas model 2 allows for a third bin, dN/dS>1; statistical tests compare models 1 & 2 to assess if there are sites with dN/dS significantly greater than 1. Similarly, model 7 includes many dN/dS bins<1, model 8a includes a bin = 1, and model 8 includes a bin with dN/dS>1. P-values were obtained by comparing the lnL values between M0 v. 0a, M1 v. 2, M7 v. 8, or M8a v. 8. We performed FUBAR using the default universal genetic code. Residues of positive selection were defined as Bayes Emperical Bayes posterior probability >0.95 (CODEML) or Bayesian based posterior probability >0.9 (FUBAR). For segments that had residues identified as positively selected by PAML, we then tested the robustness of these predictions by rerunning PAML with varying seeded omega (0.4 or 1 or 1.5) and compared multiple codon frequency models (1/61 or F3X4). See Sup Table 3 and Sup Data 1 for more detailed CODEML and FUBAR statistics and results.

### Identification of non-synonymous variants in mouse and rat strains

We examined caspase variants in a set of laboratory inbred mouse and rat strains from the Sanger mouse genome project (Lilue et al. 2018) and the rat Heterogeneous Stock founder strains (Hansen and Spuhler 1984). For the rat strains, we used the Rat Genome Database Variant Visualizer tool to identify non-synonymous variants (Laulederkind et al. 2023). For the mouse strains, we did the same thing using the Mouse strain assembly hub on the UCSC genome browser (https://hgdownload.soe.ucsc.edu/hubs/mouseStrains/hubIndex.html).

## Supplemental files

**Supplemental Table 1.** Genome statistics and sources for all primate and rodent species used.

**Supplemental Table 2.** Gene presence, absence, and pseudogene calls in each species. The “Gene Presence” tab shows for each gene in each primate and rodent species whether we recovered a sequence from the caspase-1 locus that aligned well with orthologs (“align”) and whether we recovered a blast hit to a sequenced or predicted transcript. 1 = present; 0 = pseudogene; -1 = absent; * indicates genes with partial in-frame sequences. The “Pseudogene Evidence” tab lists the SNPs, deletions, and frameshifts in each gene that led us to call it a pseudogene.

**Supplemental Table 3.** Summary of selection analyses. The “Species per Gene Analysis” tab indicates which species were included in each alignment analyzed. The “GARD breakpoints” tab lists the N-terminal and C-terminal end of each GARD breakpoint, according to the human or house mouse amino acid coordinates. The “PAML Model Comparisons” tab lists the dN/dS and log likelihood values, and p-values obtained for each model and statistical test for each caspase recombination segment. M0 vs. M0a tests for dN/dS not equal to 1 across the whole gene (i.e. deviation from drift). The other comparisons (M1 vs. M2, M7 vs. M8, M8a vs. M8) are each tests for positive selection, i.e. dN/dS>1 with alternative statistical models. All p-values are compared to a Bonferroni corrected value of 3.33E-03. The “FUBAR Summary” tab lists the number of positively selected sites and the highest Bayes factor per segment.

**Supplemental Data 1.** Alignments and full results of GARD, CODEML, and FUBAR analyses. We provide the alignments and raw data to maximize reproducibility and data transparency.

## Acknowledgements

We thank Matt Daugherty, Liz Fay, Janet Young, Patrick Mitchell, Mahtab Moayeri, Cammie Lesser, Sunny Shin, members of the Levin lab, and members of the Pitt Molecular Evolution discussion group for their helpful input on the project and manuscript. T. Levin and M. Holland were supported in this work by a grant from the National Institutes of Health (R35GM150681).

## Notes

### Competing Interest Statement

The authors have declared no competing interest.

### Summary of Updates

Added a section on caspase variants present in mouse and rodent lines, as well as analyses using basal-branching primates. We also edited for clarity.

