## Supplemental Data 1 for "Evolutionary dynamics of pro-inflammatory caspases in primates and rodents": SupData1_README.rtf

File description:
1) The figure_alignments_trees folder contains the alignments and tree files used to create Figures 1 and 3

2) The annotated_loci folder contains the extracted annotated caspase loci of primates and rodents, including exon annotations for each gene. The individual files are in Genbank format and include both sequence and annotation information.

3) The selection_analyses folder contains the alignment inputs and raw output files for each of our analyses of selection for each caspase gene.
Within selection_analyses, each gene folder contains:
1. Species tree based PAML analysis
	Alignment segments based on the GARD breakpoints
	PAML output files
		a. NS = models 0/1/2/7/8
		b. Model 8a
		c. Model 0a
		d. Varying codonFreq and starting omega
2. Segment tree based PAML analysis
	Alignment segments based on the GARD breakpoints
	PAML output files
		a. NS = models 0/1/2/7/8
		b. Model 8a
		c. Model 0a
		d. Varying codonFreq and starting omega
3. Datamonkey analyses
	GARD .json output of breakpoints
	FUBAR .csv results of each alignment segment
